# A requirement for neutrophil glycosaminoglycans in chemokine:receptor interactions is revealed by the streptococcal protease SpyCEP

**DOI:** 10.1101/461418

**Authors:** Jenny Goldblatt, Richard Ashley Lawrenson, Luke Muir, Saloni Dattani, Tomoko Tsuchiya, Shiro Kanegasaki, Shiranee Sriskandan, James Edward Pease

## Abstract

To evade the immune system, the lethal human pathogen *Streptococcus pyogenes* produces SpyCEP, an enzyme that cleaves the C-terminal *α*-helix of CXCL8, resulting in markedly impaired recruitment of neutrophils to sites of invasive infection. The basis for chemokine inactivation by SpyCEP is, however, poorly understood, as the core domain of CXCL8 known to interact with CXCL8 receptors is unaffected by enzymatic cleavage.

We examined the *in vitro* migration of human neutrophils and observed that their ability to efficiently navigate a CXCL8 gradient was compromised following CXCL8 cleavage by SpyCEP. SpyCEP-mediated cleavage of CXCL8 also impaired CXCL8-induced migration of transfectants expressing the human chemokine receptors CXCR1 or CXCR2. Despite possessing an intact N-terminus and preserved disulphide bonds, SpyCEP-cleaved CXCL8 had impaired binding to both CXCR1 and CXCR2, pointing to a requirement for the C terminal *α*-helix. SpyCEP-cleaved CXCL8 had similarly impaired binding to the glycosaminoglycan heparin. Enzymatic removal of neutrophil glycosaminoglycans was observed to ablate neutrophil navigation of a CXCL8 gradient, whilst navigation of an fMLP gradient remained largely intact.

We conclude therefore, that SpyCEP cleavage of CXCL8 results in chemokine inactivation due to a requirement for glycosaminoglycan binding in productive chemokine:receptor interactions. This may inform strategies to inhibit the activity of SpyCEP, but may also influence future approaches to inhibit unwanted chemokine-induced inflammation.

## Introduction

Chemokines and their receptors form part of a complex network, noted for their roles in positioning leukocytes and other cells via the process of chemotaxis or directed migration (1, 2). Neutrophils play a prominent part in responses of the innate immune system and are guided to sites of microbial infection by members of the ELR^+^ subgroup of CXC chemokines, which contain a Glutamate-Leucine-Arginine motif at their N-terminus. Chief amongst the ELR^+^ chemokines is CXCL8/Interleukin-8 which interacts with two principal receptors on the neutrophil surface known as CXCR1 (3) and CXCR2 (4). CXCL8 is expressed and secreted by tissue macrophages and other cells, for example, epithelial cells following bacterial infection (5) and serves to recruit neutrophils from the circulation to deal with the invading pathogen. Mice deficient in CXCR2, the major receptor for murine ELR^+^ chemokines such as KC and MIP-2, exhibit profound defects in neutrophil emigration to sites of both microbial (6) and sterile-induced inflammation supporting the notion that gradients of ELR^+^ chemokines direct the chemotaxis of neutrophils *in vivo*. In a variety of inflammatory diseases, inadvertent or overexpression of CXCL8 has been associated with pathological consequences and consequently, much effort has been put into the discovery of small molecule antagonists of CXCL8 receptors (9).

The importance of chemokines in coordinating leukocyte migration in host defense has not escaped the attention of microbes. Several pathogens have evolved ways to subvert the chemokine system and hence evade clearance by leukocytes. These include the synthesis of chemokine binding proteins by poxviruses which neutralize the *in vivo* activity of chemokines (10, 11) and the production of enzymes by hookworms which specifically degrade chemokines involved in eosinophil recruitment (12). *Streptococcus pyogenes* (group A streptococcus or GAS) is known to cause a spectrum of infections, ranging from pharyngitis and impetigo to more invasive life-threatening diseases, such as necrotizing fasciitis, which is characterized by a marked and paradoxical paucity of neutrophil recruitment at sites of severe infection and heavy bacterial growth (13). Invasive *S. pyogenes* infection is associated with the upregulation of several genes encoding virulence factors, amongst which the gene *cepA*/SpyCEP is of particular interest (14). The *cepA* gene encodes for a protein known as *Streptococcus pyogenes* cell envelope protease (SpyCEP) which specifically cleaves CXCL8 within the C-terminal *α*-helix, resulting in truncation of CXCL8 by 13 amino acids. The prominence of SpyCEP among GAS virulence factors has been ably demonstrated by loss-of-function and gain-of-function analyses (15, 16). SpyCEP-deficient strains have been found to be readily cleared in murine models of necrotizing fasciitis (15–17) whilst, conversely, heterologous expression of SpyCEP in *Lactococcus lactis* reproduced many of the features of severe necrotizing *S. pyogenes* infection in a hitherto avirulent bacterium, notably an inability to be cleared and an ability to disseminate to other organs (15). Immunity to SpyCEP confers additional protection in more virulent models of murine infection (18)

Of interest is the mechanism of action of SpyCEP, namely why C-terminal truncation of CXCL8 should result in a reduction of biological activity. Current models of chemokine:receptor activation conform to a two-step model in which firstly the chemokine receptor N-terminus interacts with the chemokine core domain (chemokine recognition site 1, CRS1), tethering and orientating the chemokine so that secondly, its N-terminus can interact with the receptor ligand-binding pocket (chemokine recognition site 2, CRS2) (19). In agreement with this model, early structure:function studies of CXCL8 and the receptors CXCR1 and CXCR2 suggested major roles for the CXCL8 N-terminus in receptor activation following ligand binding (20–22) and for the receptor N-termini in ligand recognition (23, 24). In contrast, the C-terminus of CXCL8 has been implicated in binding to glycosaminoglycans (GAGs) via a cluster of lysine residues (25, 26). Translocation of CXCL8 to the luminal surface of the endothelium is known to require interactions between the chemokine C-terminus and endothelial GAGs (27). Binding to GAGs on the surface of vascular endothelial cells also allows chemokines to form stable or haptotactic gradients, encountered by leukocytes rolling along the surface under the control of endothelial-expressed selectins (28). GAG binding is crucial for the *in vivo* activity of several chemokines, notably CCL5. Mutant CCL5 molecules that are unable to bind to GAGs have been shown to retain the ability to induce leukocyte chemotaxis *in vitro*, but not leukocyte recruitment *in vivo* when introduced into the peritoneum of mice (29). Although reduced interaction with endothelial GAGs may explain some of the effects of SpyCEP observed *in vivo*, it cannot explain the wider effects of SpyCEP on neutrophils that are evident *in vitro*, such such as reduced CD62L shedding or chemotaxis (13). If N-terminal mediated receptor binding of SpyCEP-truncated CXCL8 were preserved, and this were independent of GAG binding, some CXCR1 and CXCR2 function might be expected to be preserved.

In this study we examined the effects of SpyCEP cleavage upon neutrophil migration and the interaction of CXCL8 with the receptors CXCR1 and CXCR2 and with cell surface GAGs. We used a combination of cell transfectants and freshly isolated human neutrophils to examine different aspects of these processes. We highlight a previously unappreciated role for the C-terminal *α*-helix of CXCL8 in chemokine receptor binding and signaling, that is explained by a requirement for GAG binding. We suggest that the impaired recruitment of neutrophils during severe *S. pyogenes* infections results from not only an inability to generate a trans-endothelial chemokine gradient, but also an inability of neutrophils to respond to chemokines that are cleaved by SpyCEP.

## Materials and Methods

### Materials

All materials were obtained from Thermo Fisher Scientific (Renfrew, UK) unless otherwise stated. Recombinant human CXCL8 was obtained from Bio-Techne (Abingdon, UK) and was purchased in both the 72 amino acid (CXCL8^1-72^) and 77 amino acid (CXCL8^1-77^) forms. All experiments were performed with the 72 amino acid form of CXCL8 unless stated otherwise. Other recombinant chemokines were from Peprotech EC (London, UK). *N*-Formyl-Met-Leu-Phe (fMLP) was from Bio-Techne. The glycanase cocktail of heparinase I, heparinase II and chondroitinase ABC was purchased from Sigma-Aldrich (Poole, UK).

### Cell culture and Transfection

The mouse pre-B cell line L1.2 was maintained and transfected with plasmids by electroporation as previously described (30). The plasmid vector pcDNA3 encoding HA-tagged variants of human CXCR1 and CXCR2 were purchased from the cDNA Resource Center (Bloomsburg University, PA, USA). Four hours following transfection, cultures were supplemented with Sodium Butyrate (Sigma-Aldrich) at a final concentration of 10mM. Overnight culture in the presence of sodium butyrate enhances the transient expression of chemokine receptors in this system. Expression of HA-tagged CXCR1 or CXCR2 was confirmed prior to experimentation (data not shown) by the use of an anti-HA monoclonal and flow cytometry analysis by FACS Calibur (BD Bioscience, Oxford, UK) as previously described (30).

Human neutrophils were isolated from whole blood obtained from a subcollection of the Imperial College Tissue Bank, taken from informed, consenting healthy normal subjects. Neutrophils were freshly isolated by negative selection using the MACSxpress neutrophil isolation kits according to the manufacturer’s instructions (Miltenyi Biotec Ltd, Woking, UK) followed by a single RBC lysis step (using hypo / hypertonic solutions).

### Chemokine cleavage by SpyCEP

Emm81 *S. pyogenes* strain H292 has been previously defined as a high SpyCEP producing strain (13, 14) and was used as a source of SpyCEP with a molecular weight of approximately 160kDa. H292 was grown for 16 hours at 37 °C in an atmosphere of 5% CO_2_ in RPMI (ThemoFisher Scientific) and the supernatant retained. Culture supernatants were centrifuged at 2500g for 10 minutes at 4 °C to pellet bacteria, following which they were passed through a 0.2 µm filter (VWR, Lutterworth, UK), split into aliquots stored at − 20 °C. GAS strain H575 is an isogenic mutant of H292, in which the majority of *cepA* has been deleted, leading to production of an inactive truncated N-terminal SpyCEP fragment of approximately 40 kDa (15). Supernatants from this strain were produced in an identical fashion as a negative control for SpyCEP cleavage of chemokines. Chemokine cleavage assays were carried out by digesting a known amount of recombinant CXCL8 at 37 °C for 16 hours in supernatant from the H292 strain (+ SpyCEP) or H575 (-SpyCEP) GAS strains after which it was diluted in assay buffer (RPMI + 0.1% BSA) to give the required concentration of chemokine.

### Chemotaxis Assays

Dilutions of CXCL8 treated with either the H575 or H292 supernatants were made in assay buffer (RPMI + 0.1 % BSA). Migration towards CXCL8 was assessed using L1.2 cell transfectants expressing HACXCR1 or HA-CXCR2 in modified Boyden Chamber assays as previously described (30) using ChemoTX chambers with a 5 µm pore size (Neuroprobe Inc., Gaithersburg, MD).

For the real time analysis of migrating neutrophils, a 12-channel TAXIScan was employed (31) and used according to the manufacturer’s protocol (Effector Cell Institute, Tokyo. Japan). 1 µl of a suspension containing 5 × 10^5^ neutrophils/ml was loaded into each chamber and following alignment of the cells at one end of the terrace, 1 µl of 100nM CXCL8 or 100nM fMLP was added to the opposing end of the terrace (260 µm away) and cells were allowed to migrate along the ensuing chemoattractant gradient for 1hr at room temperature. Sequential image data were captured every minute as individual jpegs which were subsequently processed with ImageJ (National Institutes of Health), equipped with the manual tracking (Fabrice Cordelieres, Institut Curie, Orsay (France) and chemotaxis tool plugins (Ibidi, Martinsried, Germany).

Individual experiments consisted of duplicate conditions for each chemokine and data illustrated are collated from an equal number of experiments as highlighted in the figure legend. The total numbers of cells tracked under each condition are shown in the top right hand corner of each plot. For each individual cell, a variety of parameters were generated via the chemotaxis tool plugin, namely the accumulated distance travelled by each cell, its velocity, its directionality and its forward migration index parallel to the gradient (FMI^||^). Directionality is defined as the ratio of Euclidian distance:accumulated distance travelled. A value of 1 represents migration in a perfectly straight line. The FMI^||^ is defined as the distance travelled by the cell in the y axis (i.e. along the chemokine gradient) divided by the accumulated distance it travelled (32).

In experiments employing a glycanase cocktail of heparinase I, heparinase II and chondroitinase ABC, neutrophils were incubated with agitation with one unit/ml of enzyme for 30 minutes at 37 °C as previously described (33) before being washed once in phosphate buffered saline prior to their immediate use in TAXIScan assays.

### Ligand Binding assays

Competitive binding assays were carried out with L1.2 cells expressing HA-CXCR1 or HACXCR2 using a previously described protocol (30). ^125^ICXCL8 was used as a radiolabel and was purchased from Perkin Elmer Life Sciences (Boston, MA). Competition was with increasing concentrations of unlabelled CXCL8 incubated with either H292 or H575 culture supernatants and subsequently treated with Pefabloc SC (Roche, Lewes, UK), a serine protease inhibitor which inactivates SpyCEP (13). This was a precaution against digestion of the ^125^I-CXCL8 tracer.

### Statistical Analysis

Statistical analyses were carried out using Prism 6 (GraphPad Software, La Jolla, CA) and the tests are noted in the figure legends. * = P<0.05, **= P<0.01, ***= P<0.001 and ****= P<0.0001.

## Results

### SpyCEP has broad spectrum activity for ELR^+^ CXC chemokines

To ascertain the specificity of SpyCEP for neutrophil-recruiting CXC chemokines, a selection of recombinant chemokines with varying degrees of homology (Figure 1A) were incubated overnight with washed wild type S. pyogenes cells (H292) (14) or the isogenic *cepA* mutant (H575) (15). Incubation with wild type *S. pyogenes* cells evidently resulted in the cleavage of all ELR^+^ CXC chemokines as deduced by SDS-PAGE analysis (Figure 1B), with a notable reduction in their molecular weight. In contrast, the non-ELR^+^ chemokines included as controls remained intact (CXCL4, CXCL9 and CXCL10). Incubation with control, SpyCEPnegative (H575) cells resulted in no detectable cleavage of any of the chemokines examined.

**Figure. 1.**
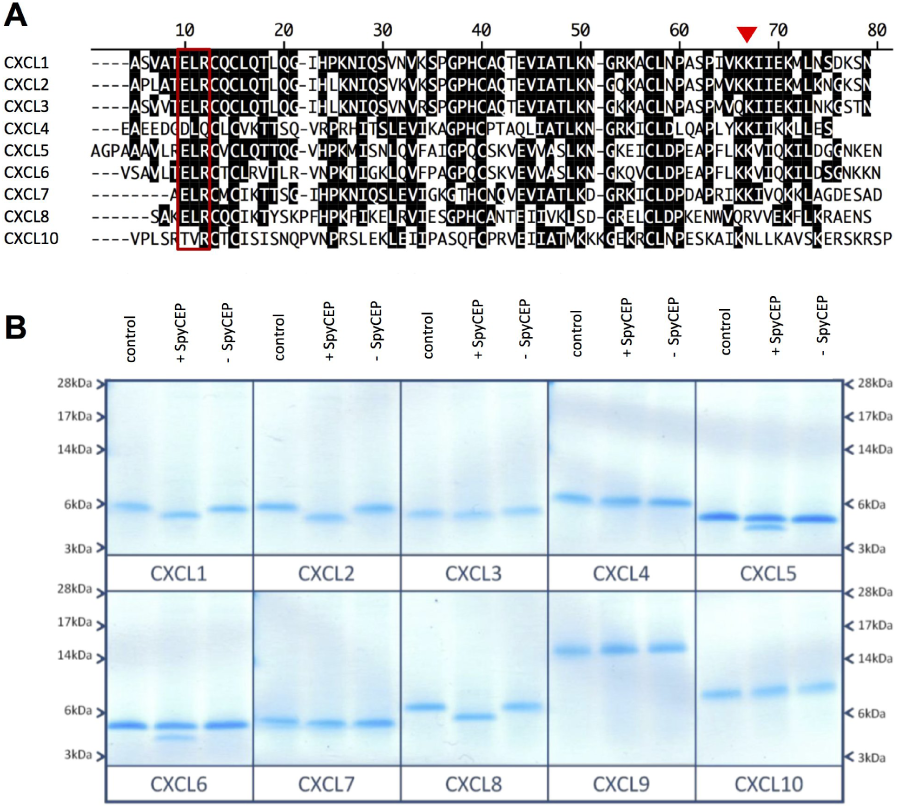
SpyCEP has broad spectrum activity for ELR^+^ CXC chemokines. Panel A shows an alignment of a selection of mature human CXC chemokine sequences. The ELR motif shared by several of the chemokines is boxed and the previously determined site of cleavage of CXCL8 by SpyCEP denoted by a red triangle (13). Panel B shows the effects of SpyCEP cleavage on chemokine. 500ng of chemokine was incubated with 2 µl of washed wild type GAS cells (H292) or an isogenic *cepA* mutant (H575). Cleavage was allowed to proceed for 18hr at 37 °C after which the proteins were separated by SLS-PAGE and stained with colloidal blue (Panel B)

### Truncation of CXCL8 by SpyCEP results in impaired navigation of a chemokine gradient

The TAXIScan instrument was employed to assess the real-time migration of freshly isolated neutrophils along a gradient of intact CXCL8 or SpyCEP-cleaved CXCL8 over a 1 hr period. Intact CXCL8 was seen to elicit rapid neutrophil migration with the majority of cells traversing the terrace in under 30 minutes in a direct fashion (Figure 2A, Supplementary video 1). SpyCEP-cleaved CXCL8 induced the migration of far fewer neutrophils which lacked focus and appeared hesitant, making more turns and correspondingly taking much longer to traverse the terrace (Figure 2B, Supplementary video 2). Basal neutrophil migration in the absence of stimulus was minimal (Fig. 2C). Our observations were confirmed by single-cell tracking analysis with SpyCEP-cleaved CXCL8 inducing significantly slower migration than intact CXCL8 (Fig. 2D), accompanied by a significantly lower directionality component (Fig. 2E). When the forward migration indices parallel to the chemokine gradient were calculated (FMI^||^), SpyCEP-cleavage of CXCL8 was also seen to significantly impair the extent of migration compared to intact CXCL8 (Fig. 1F). Thus, SpyCEP cleavage of CXCL8 significantly impairs migration along the chemokine gradient, resulting in slower, less directed neutrophil responses. Similar data sets were also obtained with the N-terminally extended form of CXCL8 (CXCL8^1-77^) which is expressed by endothelial cells (Figure 3A-F). Thus, SpyCEP is able to cleave both biologically active forms of CXCL8, rendering them less efficacious in terms of leukocyte recruitment.

**Figure. 2.**
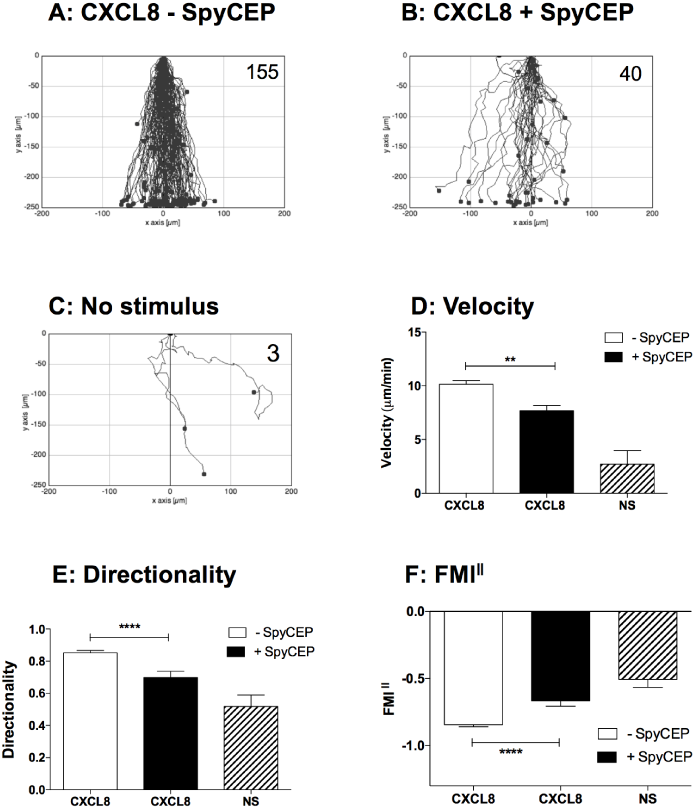
SpyCEP cleavage of CXCL8 results in significantly impaired migration of neutrophils as visualized in realtime by TAXIScan. Panels A-C show the collated tracks of individual migrating neutrophils (duplicate conditions) pooled from three independent experiments using different donors. Gradients of H575 or H292 treated CXCL8 were established in panels A and B respectively, while panel C shows the lack of neutrophil migration in the absence of chemokine. The total number of tracked cells from all three experiments is shown in the top right hand corner. Panels D-F show significant differences in velocity, directionality and forward migration index parallel to the gradient (FMI^||^) of the data in panels A-C following tracking analysis. Error bars represent the SEM. Statistical significances between CXCL8 treatments were determined by one-way ANOVA with Tukey’s’ post-test

**Figure. 3.**
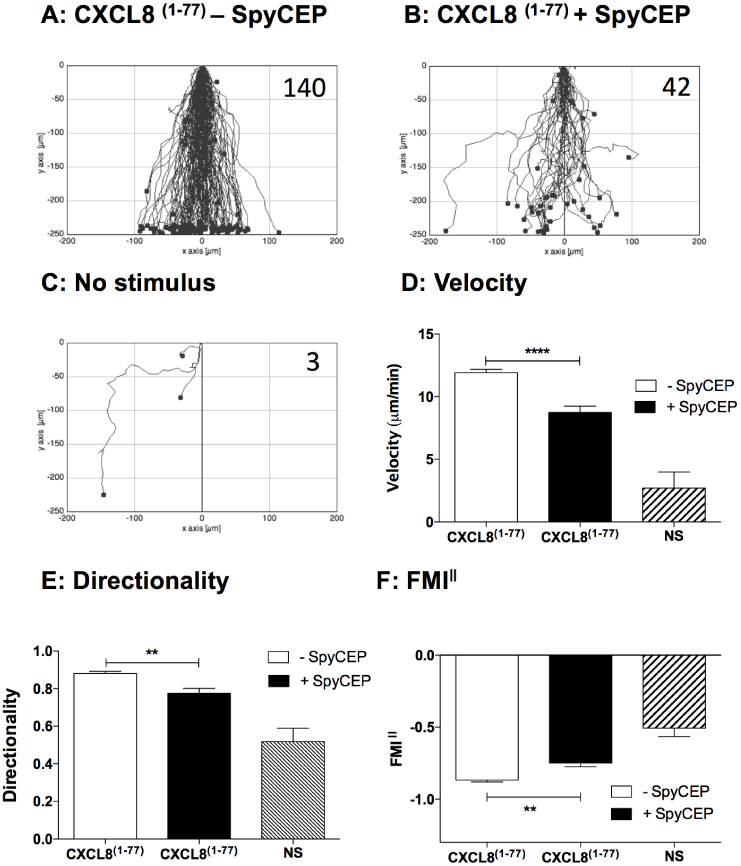
SpyCEP cleavage of the extended form of CXCL8 (CXCL8^1-77^) results in significantly impaired migration of neutrophils. Panels A-C show the collated tracks of individual migrating neutrophils (duplicate conditions) pooled from three independent experiments using different donors. Gradients of H575 or H292 treated CXCL8^1-77^ were established in panels A and B respectively, while panel C shows the lack of neutrophil migration in the absence of chemokine. The total number of tracked cells from all three experiments is shown in the top right hand corner. Panels D-F show significant differences in velocity, directionality and FMI^||^of the data in panels A-C following tracking analysis. Error bars represent the SEM. Statistical significances between CXCL8^1-77^ treatments were determined by one-way ANOVA with Tukey’s’ post-test.

### Truncation of CXCL8 by SpyCEP results in reduced activation and binding to CXCR1 and CXCR2

Previous reports have detailed the effects of CXCL8 truncation by SpyCEP on neutrophil function (13), but have not determined the relative contributions of CXCR1 and CXCR2 signaling. We set out to address this using constructs encoding N-terminal HA-tagged variants of CXCR1 and CXCR2. These were transiently expressed at high levels in the mouse pre-B cell L1.2 according to published protocols (34), allowing ligand binding assays and chemotaxis assays to be performed.

We firstly compared the ability of either SpyCEP-cleaved or intact CXCL8 to induce migration of CXCR1 and CXCR2 transfectants in modified Boyden chamber assays (Figure 4). Intact CXCL8 was efficacious in recruiting both CXCR1 and CXCR2 transfectants, with optimal migration seen in the 110 nM concentration range (Fig. 4A and B). In contrast, cleavage of CXCL8 with SpyCEP resulted in a significant reduction in both potency and efficacy compared with intact CXCL8. Previous mass spectrometry analysis confirms that the N–terminal domain and disulphide bridges within CXCL8 remain intact following SpyCEP cleavage (13), retaining the chemokine fold required for receptor binding. However, the reduced ability of SpyCEP-cleaved CXCL8 to provide a chemotactic gradient for neutrophils or for transfectants raised the possibility that cleaved CXCL8 might be unable to bind to its cognate receptors CXCR1 and CXCR2. Cleaved or uncleaved CXCL8 samples were diluted in assay buffer and activity in competitive binding assays was evaluated using CXCR1 and CXCR2 transfectants. As expected, intact CXCL8 that had been incubated with the control (H575) supernatant readily displaced ^125^I-CXCL8 from both CXCR1 and CXCR2 transfectants at nanomolar concentrations (IC_50_ values of 8.1nM and 8.7nM at CXCR1 and CXCR2 respectively, Fig 4C and D). In contrast, cleaved CXCL8 that had been incubated with SpyCEP was unable to displace 50% of the ^125^I-CXCL8 from either receptor, even when used at a 1000-fold greater concentration than the labelled ligand, consistent with the reduced ability to signal through CXCR1 or CXCR2 observed in the migration assays. Thus, we conclude that it is the loss of the terminal 13 residues alone that adversely affects the binding of CXCL8 to both CXCR1 and CXCR2 and subsequent receptor activation.

**Figure. 4.**
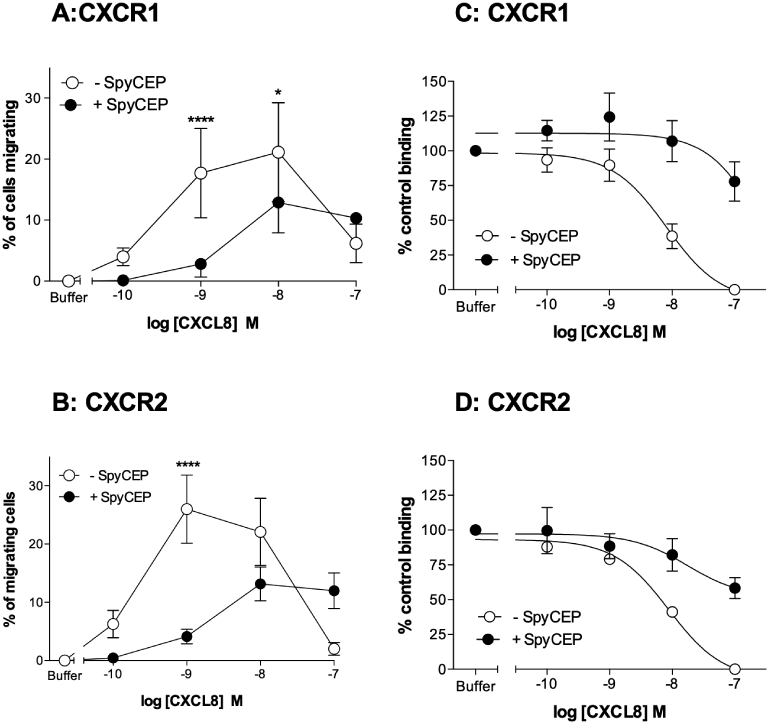
SpyCEP cleavage of CXCL8 inhibits binding and activation of CXCR1 and CXCR2. Panels A and B show the ability of CXCL8 to compete for the binding of 125I-CXCL8 to CXCR1 or CXCR2 transfectants following incubation with supernatants from either the H292 or H575 GAS strains (n=6).Panels C and D show the activities of identically treated CXCL8 in inducing the migration of CXCR1 or CXCR2 transfectants (n=6). Error bars represent the mean ± SEM.

### Truncation of CXCL8 by SpyCEP results in reduced glycosaminoglycan binding

Chemokines have been shown to adhere with micromolar affinity to glycosaminoglycans (GAGs) which is essential for chemokine presentation on endothelial cells and the recruitment of leukocytes into tissues *in vivo* (35). The GAG binding domain of CXCL8 was shown by Kuschert *et al* to be comprised of five basic amino acids, namely K20 (within a region known as the proximal loop) and R60, K64, K67, and R68 in the C-terminal *α*-helix (36). Since truncation by SpyCEP at Q59 would remove four of these residues, we hypothesized that it may have deleterious effects on GAG binding. We therefore examined the ability of CXCL8 to form oligomers on heparin sepharose beads following treatment with SpyCEP containing supernatants. In agreement with previous data from Hoogewerf and co-workers (36), increasing concentrations of intact CXCL8 lead to a corresponding increase in the proportion of CXCL8 bound to the beads (Fig 5). In contrast, SpyCEP-cleaved CXCL8 was without activity in this assay, suggesting that removal of a large proportion of the GAG binding site by SpyCEP (4 out of 5 basic residues) renders CXCL8 unable to bind to heparin.

**Figure. 5.**
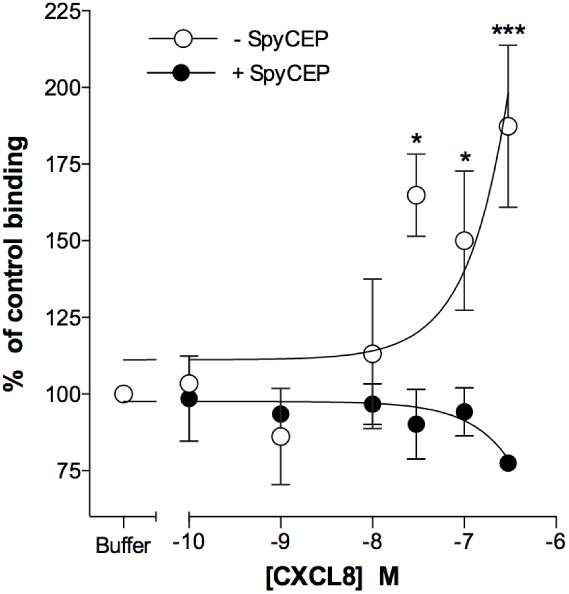
SpyCEP cleavage of CXCL8 ablates binding to heparin. Graph shows the relative abilities of H292 and H575-treated CXCL8 in binding to heparin sepharose beads. Error bars represent the mean ± SEM (n=6). Statistical significance between CXCL8 treatments was determined by two-way ANOVA with Sidak’s post-test

### Effective navigation of a CXCL8 gradient requires neutrophil GAGs

Since SpyCEP cleavage impaired both GAG binding and neutrophil recruitment *in vitro*, we postulated that binding to GAGs on the neutrophil surface is a key step in the productive activation of neutrophil CXCL8 receptors. To test this postulate, we incubated neutrophils at 37°C in a glycanase cocktail containing heparanase and chondroitanase, since both heparan sulfate and chondroitin sulfate-decorated GAGs have been shown to bind CXCL8 *in vitro* (37). After incubation, neutrophils were washed once in buffer then assessed via TAXIScan for their ability to migrate along gradients of the tripeptide chemoattractant fMLP (deemed too small and uncharged to effectively bind to GAGs) or gradients of intact CXCL8. Incubation of neutrophils in buffer alone resulted in elevated basal migration which appeared to be without direction in the absence of a stimulus (Fig. 6A). The introduction of gradients of fMLP or CXCL8 induced obvious directional migration (Fig. 6B and C). Glycanase treatment resulted in an evident reduction in basal migration (Fig. 6D), although responses to fMLP remained intact (Fig. 6E). In contrast, migratory responses to CXCL8 were abolished by glycanase treatment (Fig. 6F). These findings were corroborated by single cell tracking analysis. Glycanase treatment of neutrophils reduced the velocity and directionality of the CXCL8 responses so far as they were indistinguishable from those seen in the absence of a stimulus (Fig. 6G and H). In contrast, despite glycanase treatment, fMLP-induced neutrophil migration remained significantly faster and more directional than that seen in the absence of a stimulus. Analysis of the FMI^||^ indices further clarified these findings, with glycanase treatment of neutrophils seen to ablate migration along the CXCL8 gradient, whilst migration along the fMLP gradient remained significantly greater than that seen in the absence of a stimulus (Fig 6I). Thus, we conclude that effective navigation of a CXCL8 gradient requires intact GAGs on the neutrophil cell surface, providing a potential explanation for the inactivating effects of C-terminal cleavage of CXCL8 by SpyCEP.

**Figure. 6.**
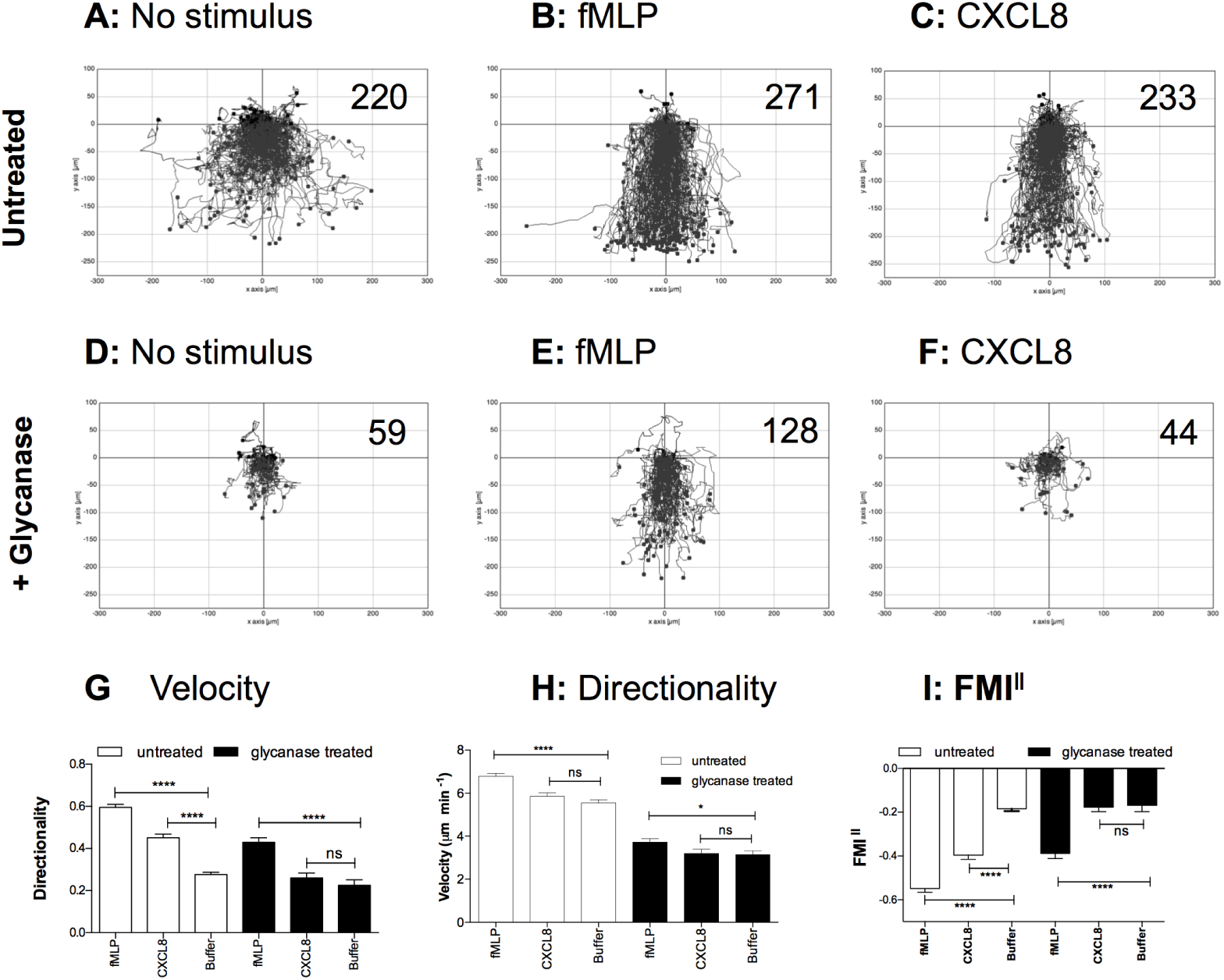
Removal of neutrophil GAGs specifically ablates migration along a CXCL8 gradient. Panels A-F show the collated tracks of individual migrating neutrophils (duplicate conditions) pooled from three independent experiments using different donors. In panels A-C, neutrophils were incubated in buffer alone prior to migration, while in panels D-F, cells were treated with a glycanase cocktail. Gradients of fMLP, CXCL8 were established where noted, with the total number of tracked cells shown in the top right hand corner. Panels G-I show significant differences in velocity, directionality and FMI^||^ of the data in panels A-C following tracking analysis. Error bars represent the SEM. Statistical significances between conditions were determined by one-way ANOVA with Tukey’s’ post-test.

## Discussion

The paucity of neutrophils in severe necrotising *S. pyogenes* infection has been directly attributed to the activity of SpyCEP, expression of which is upregulated in invasive isolates (15, 16). It was previously hypothesized that the cleavage of the CXCL8 C-terminal *α*-helix by SpyCEP resulted in abrogation of trans-endothelial chemokine gradients through the inability of cleaved CXCL8 to translocate to the luminal endothelial surface (13, 27). While this may indeed be important, it cannot explain the inactivation of CXCL8 that we and others have observed in *in vitro* assays with neutrophils. Here, we have demonstrated that SpyCEP cleavage of CXCL8 renders the chemokine unable to bind productively to its cognate receptors, providing an explanation for the observed inactivation. The broad activity of SpyCEP for a range of ELR^+^ CXC chemokines, coupled with the ability of *S. pyogenes* to cleave both C3a and C5a (38) demonstrates a potential for this lethal pathogen to abrogate major components of the human neutrophil chemoattractant repertoire. We also determined that SpyCEP cleavage of CXCL8 abolished the ability of the chemokine to bind to heparin. These observations generated the hypothesis that interaction with GAGs might be central to activation of the receptors CXCR1 and CXCR2 by CXCL8. This is supported by our demonstration that the removal of neutrophil surface GAGs by glycanases can abolish the ability of freshly isolated neutrophils to respond to CXCL8, but leave responses to fMLP broadly intact. This is reminiscent of a previous report by Hoogewerf and colleagues in which binding of ^125^I-CXCL8 to CXCR1^+^ transfectants was significantly reduced following treatment with a mixture of glycanases (36). The findings are also in keeping with an earlier report in which significant reductions in chemotaxis and receptor binding were observed with a synthetic variant of CXCL8 truncated at position 60 (23).

Placing this information in the context of what we already know about chemokine: endothelial GAG interactions (35), we propose that in the context of migration, the neutrophil glycocalyx serves several purposes. The first is the sequestration of chemokines on the neutrophil surface (both monomer and higher order species), in effect ‘sampling’ the gradient at the leading edge of the migrating leukocyte. This sequestration leads to a productive interaction with the chemokine receptors which can be envisaged as increasing the local concentration of chemokine in the vicinity of the receptors, or even a physical *cis*-presentation of chemokine to the receptors (Fig. 7). In the absence of the gradient sampling afforded by GAGs, migration along the chemokine gradient is much less efficient. As an analogy, consider an automobile taking a bend in the road after sunset. The beams from the headlights enable the driver to sample the environment and seeing the bend, steer the vehicle appropriately with negligible loss of speed. In contrast, in the absence of headlights and reduced environmental sampling, as the driver approaches the bend it is necessary to slow down to re orientate the vehicle and safely navigate the bend, leading to less fluent movement along the road. It is interesting to note that morphological differences in neutrophils navigating gradients of fMLP or CXCL8 have been reported previously by Yamauchi and colleagues. Notably, neutrophils migrating along an fMLP gradient displayed a fan-like widely-spread lamellipodium at the leading edge with a compact body and short tail. In contrast, neutrophils navigating a gradient of CXCL8 showed a more focused lamellipodium with a longer cell body and tail (39).

**Figure. 7.**
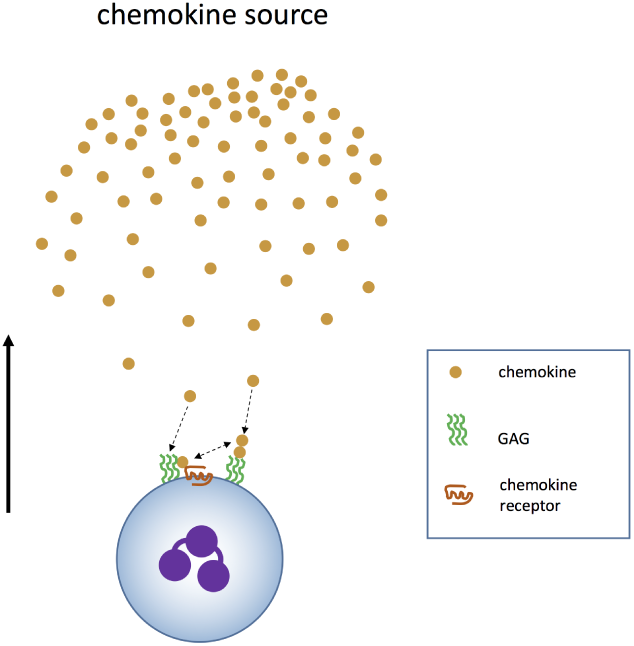
Model of enhanced gradient navigation by neutrophils due to the cooperation of chemokine receptors and GAGs. The cartoon shows a neutrophil migrating along a gradient of chemokine towards the source (direction indicated by the arrow). Cell surface GAGs encounter chemokine and bind it, enabling presentation to the chemokine receptor or increasing the local concentration of chemokine surrounding the neutrophil by assisting in chemokine oligomerization. Chemokine monomer can dissociate from the oligomer and activate the chemokine receptor (double headed arrow).

We conclude that GAGs on the surface of the neutrophil are an essential component of effective migratory responses to CXCL8 and that SpyCEP ruthlessly exploits this to render CXCL8 inactive *in vivo* (14). This raises several interesting questions. Firstly, is sampling by GAGs required for efficient navigation of other chemokine gradients by neutrophils? This question requires additional experimentation to answer conclusively. There is perhaps a clue in the broad spectrum activity of SpyCEP for all ELR^+^ chemokines (40) and Fig. 1B), although, it could be that in evolving activity against a principal neutrophil chemoattractant in CXCL8, SpyCEP has ‘unwittingly’ acquired the ability to degrade other neutrophil attractants. Secondly, are GAGs on the surface of other leukocytes critical for the navigation of chemokine gradients? Previous studies of monocytes (33) and T-cells (41) have shown that glycanase treatment results in reduced intracellular signalling in response to the chemokines CXCL4 and CCL5, although these studies were not extended to analyses of cell migration. Finally, can we learn lessons from SpyCEP in terms of targeting the interactions of GAGs with chemokines for therapeutic benefits? Mutants of CXCL8 which are likely to bind to SpyCEP but have reduced activity at CXCR1 and CXCR2 have been described by others and may be a useful starting point for drug discovery (42, 43). Provided such molecules clear the typical hurdles of bioavailability and target occupancy that have beset many small molecule chemokine receptor antagonists (44), they may present themselves as candidate SpyCEP inhibitors with potential for the adjuvant management of invasive group A streptococcal infections.

## Supporting information

